# Orbitofrontal and Thalamic Influences on Striatal Involvement in Human Reversal Learning

**DOI:** 10.1101/246371

**Authors:** Tiffany Bell, Angela Langdon, Michael Lindner, William Lloyd, Anastasia Christakou

**Author notes:** Corresponding author: Dr Anastasia Christakou, School of Psychology and Clinical Language Sciences, University of Reading, Reading RG6 6AL.

## Abstract

Cognitive flexibility is crucial for adaptation and is disrupted in neuropsychiatric disorders and psychopathology. Human studies of flexibility using reversal learning tasks typically contrast error trials before and after reversal, which provides little information about the mechanisms that support learning and expressing a new response. However, animal studies suggest a specific role in this latter process for the connections between the dorsal striatum and the centromedian parafascicular (CM-Pf) thalamus, a system which may recruit the striatal cholinergic interneurons, but which is not well understood in humans. This study investigated the role of this system in human probabilistic reversal learning, specifically with respect to learning a new response strategy, contrasting its function to that of the better understood orbitoftontal-striatal systems. Using psychophysiological interaction (PPI) analysis of functional magnetic resonance imaging (fMRI) data we show that connectivity between the striatum and both the lateral orbitofrontal cortex (lOFC) and CM-Pf pathways increased during reversal, but not initial learning. However, while the strength of lOFC-striatal connectivity was associated with the speed of the reversal, the strength of CM-Pf-striatal connectivity was associated specifically with the quality of the reversal (reduced regressive errors). These findings expand our understanding of flexibility mechanisms in the human brain, bridging the gap with animal studies of this system.

## INTRODUCTION

Cognitive flexibility, altering behaviour following changes in the environment, is disrupted in neuropsychiatric disorders, including Parkinson’s disease, autism, obsessive compulsive disorder, and schizophrenia (Nilsson et al., 2015; Prado et al., 2017). Cognitive flexibility can be measured using reversal learning tasks, where a previously formed stimulus-reward association is degraded, and a different stimulus becomes relevant for guiding behaviour (Nilsson et al., 2015).

Multiple brain areas have been shown to be recruited during reversal learning. Functional magnetic resonance imaging (fMRI) shows activity related to reversal learning in frontoparietal cortex (Cools et al., 2002; Remijnse et al., 2005; D’Cruz et al., 2011; Waegeman et al., 2014), the striatum (Remijnse et al., 2005; Ghahremani et al., 2010), hippocampus (Vilà-Balló et al., 2017), and amygdala (Yaple and Yu, 2019). In particular, human studies support an emerging consensus from animal work regarding distinct roles of orbitofrontal cortical (OFC) areas, with the medial and lateral portions of the OFC shown to contribute primarily to initial versus reversal learning (Dalton et al., 2016; Morris et al., 2016; Izquierdo et al., 2017), by supporting attention versus credit assignment respectively (Rushworth et al., 2011; Noonan et al., 2017).

Clearly these brain regions support reversal learning as part of a network (Yaple and Yu, 2019), but studying the configuration of functionally meaningful cortico-subcortical networks during time-variant task performance remains a considerable challenge. The striatum in particular is known to play a key role during the reversal learning process, however this knowledge is largely due to a wealth of evidence from animal studies, while there is little neuroimaging evidence of its precise involvement in humans (Remijnse et al., 2005). Region-of-interest and multimodal approaches enable us to capitalise on insights from animal work and circumvent some of the challenges of functional neuroimaging (Morris et al., 2016).

Additionally, human studies typically use two stimuli and multiple reversals, often focusing on contrasts of activation during different error types (e.g. before versus after the reversal) (Ghahremani et al., 2010; Izquierdo et al., 2017). Although this approach can help characterise the broad inhibitory and attentional mechanisms involved, it tells us less about the mechanism through which the reversal itself is identified, and less still about the mechanism required for discovering and implementing a new behaviour.

To understand these mechanisms, animal research points to the striatal cholinergic interneuron (CIN) system, which modifies the action of corticostriatal circuitry, gating striatal-level input from the centromedian-parafascicular nucleus (CM-Pf) of the thalamus (Ellender et al., 2013; Smith et al., 2014; Yamanaka et al., 2018; Assous and Tepper, 2019). This system is of particular interest because it is implicated in pathological cognitive inflexibility, for example, in neurodegenerative disorders (Henderson et al., 2000; Smith et al., 2014) and schizophrenia (Holt et al., 1999, 2005).

The CM-Pf – striatal cholinergic system plays a role distinct to OFC which is involved in attention and credit assignement (Noonan et al., 2017). In animals, disruption of cholinergic signalling in the dorsomedial striatum increases regressive errors during reversal learning, both when CINs are ablated, and when input from the CM-Pf is impaired (Brown et al., 2010; Bradfield et al., 2013). The specificity of these effects, and the fact that this system interfaces with OFC-striatal circuitry (Stalnaker et al., 2016), together offer important insight into neurocognitive flexibility mechanisms. In humans, using functional magnetic resonance spectroscopy (fMRS), we recently demonstrated task-related changes in cholinergic activity in the dorsal striatum during uninstructed multialternative probabilistic reversal (Bell et al., 2018, 2019). Here, we ask whether CM-Pf-striatal connectivity is also implicated in human reversal learning, and – if so – what is its distinct contribution relative to the better understood OFC-striatal system.

We hypothesised that both lateral OFC-striatal and CM-Pf-striatal connectivity would be involved in reversal learning, but would support distinct aspects of performance as described above. We used the medial OFC and mediodorsal nucleus of the thalamus (MD) as specificity controls, given evidence for dissociable contributions of these regions to learning, as well as discrete connectivity patterns (described in the methods section). We used high-resolution multiband fMRI and psychophysiological interaction analysis (PPI) to test for changes in connectivity in our dyads of interest during multialternative probabilistic reversal learning.

## METHODS

### Participants

The study was approved by the University of Reading Research Ethics Committee (reference: UREC13/15). Fifty seven (57) volunteers (20 female) between the ages of 18.5 and 30.6 (mean = 22.7, SD = 3.6) were recruited by opportunity sampling from the University of Reading and surrounding areas. All participants were healthy, right handed non-smokers and written informed consent was taken prior to participation.

One participant was excluded due to technical issues during data collection. 34 participants were excluded from the analysis reported here as they did not reach the task learning criteria specified below (and as previously applied, Bell et al., 2018). Twenty two (22) participants reached criterion in both initial and reversal learning and were included in the analysis (12 female; mean age = 22.5, SD = 3.8).

### Behavioural Data

#### Learning Task

The task was a probabilistic multi-alternative reinforcement learning task with a reversal component, described previously (Bell et al., 2018). It was programmed using MATLAB (2014a, The Mathworks, Inc., Natick, MA, United States) and Psychtoolbox (Brainard, 1997).

Four card backs were displayed on a screen representing four decks of cards. Win cards resulted in +50 points and loss cards resulted in −50 points. Each deck had a different probability of generating a win card (75%, 60%, 40%, and 25%) and the probabilities were randomly assigned across the four decks for each participant. Outcomes were pseudo-randomised so that the assigned probability was true over every 20 times that deck was selected. Additionally, no more than four cards of the same result (win/loss) were presented consecutively in the 75% and 25% decks and no more than three cards of the same result in the 60% and 40% decks.

Participants chose a deck by pressing the corresponding button on a four-button box. A cumulative points total was displayed in the centre of the visual display at the end of each trial, and in the bottom right hand corner throughout the session. The trial structure, including timings is shown in Figure 1A.

**Figure 1.**
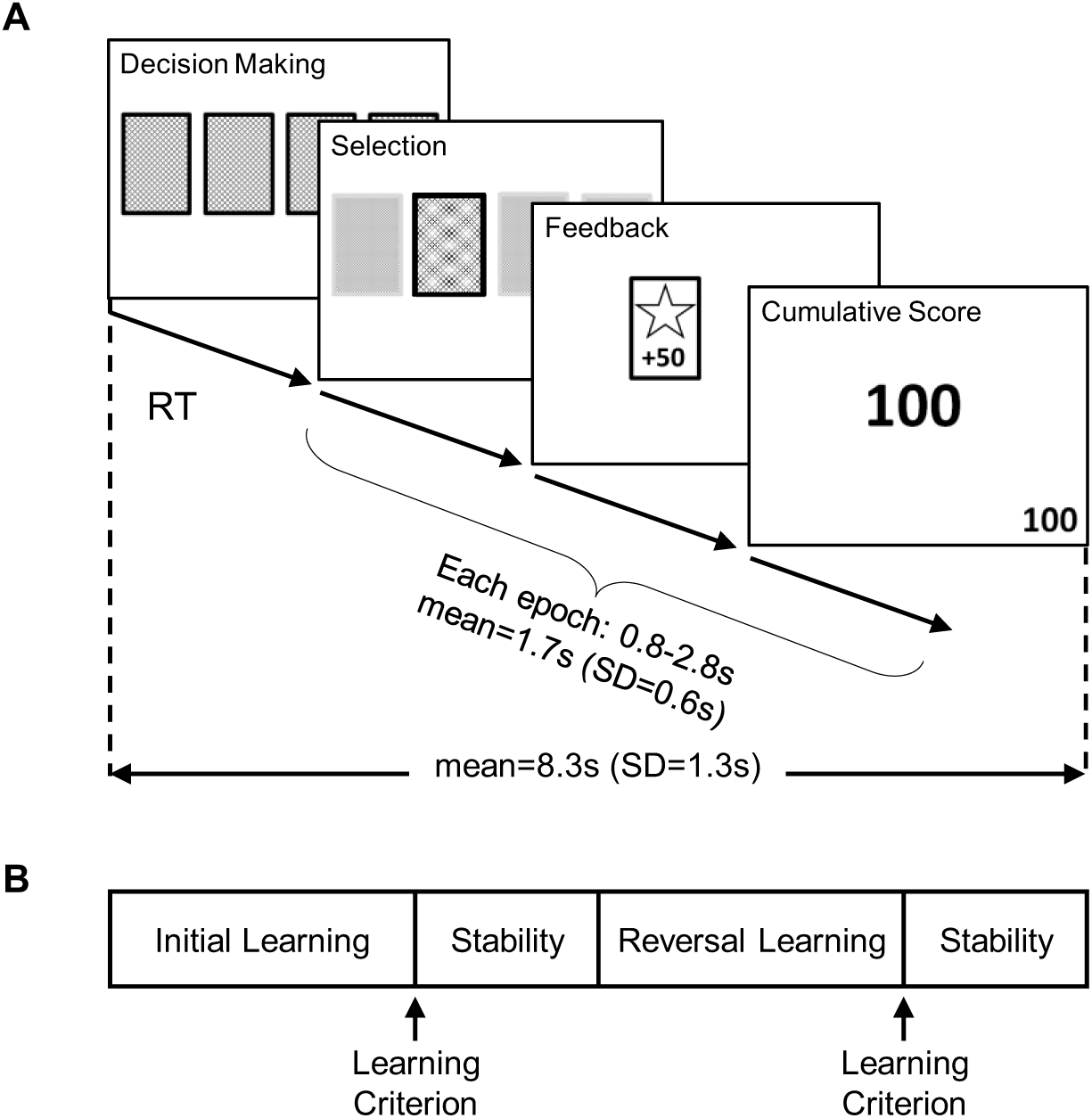
Trial and task design. A. Participants were instructed to choose between four decks of cards. Each deck had a different probability of generating winning cards (75%, 60%, 40% and 25%). Once the predetermined learning criterion had been reached, the deck probabilities were reversed so that high probability decks became low probability decks and vice versa. Participants were not informed of this in advance and were simply instructed to gain as many points as possible. The time from initial deck presentation to deck choice is the decision-making epoch referred to in the analysis. The length of time the feedback was displayed was the feedback epoch referred to in the analysis. RT = reaction time; SD = standard deviation. B. Schematic of the four task phases. Upon reaching criterion in the initial learning phase, participants completed a post criterion stability phase (lasting for 60% of the trials-to-criterion during initial learning). After this phase, the deck probabilities were reversed. Participants then completed a post-reversal learning phase and upon reaching criterion again, they completed another post criterion stability phase (lasting for 60% of the trials-to-criterion during reversal learning).

The task was split into four phases: initial learning, first stability, reversal learning and post-reversal stability (Figure 1B). As the research question focused on the reversal, we wanted to encourage behavioural stability before the reversal to reduce intra-individual noise. Therefore, a “stability phase” was included at the end of the initial learning phase. The number of trials in this phase was equal to 60% of the number of trials taken to reach criterion. Participants only moved onto the next phase if the learning criterion (selection of either of the two highest decks on at least 80% of 20 consecutive trials) was reached. Participants were given 100 trials to reach criterion in both the initial learning and reversal phase. If participants did not reach criterion in the initial learning phase, they did not experience the first stability or reversal. Following the reversal, participants were allowed 100 trials to reach criterion and enter the post-reversal stability phase, after which the task ended (Figure 1B).

Performance was measured using the number of trials taken to reach criterion during the initial learning and reversal phases. Perseverative errors were defined as the trials after reversal until the probability of selecting the previously favoured deck reached chance level (0.25), i.e. the number of trials taken to identify the reversal and switch behaviour. Regressive errors were defined as selections of the previously favoured deck after the perseverative period had ended.

#### Temporal Difference Reinforcement-Learning (TDRL) Model

We used a temporal difference reinforcement learning algorithm (Sutton and Barto, 1998) to model behaviour as a function of previous choices and rewards, as described previously (Bell et al., 2019). Briefly, a soft-max probability distribution was used to describe the probability that deck *c* was chosen on each trial *t*,

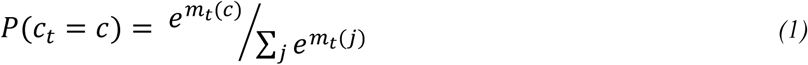

where *m*_*t*_(*c*) is the preference for the chosen deck, and *j* indexes the four available decks. To determine the preference for the chosen deck, the expected value of that deck on a given trial *V*_*t*_(*c*), was multiplied by the participant’s individual value impact parameter *β* (equivalent to the inverse temperature):

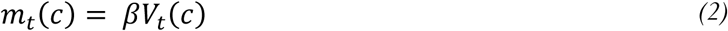

A prediction error (*pe*_*t*_) was calculated on each trial by subtracting the expected value of the chosen deck from the actual reward (a win or loss: *reward*_*t*_, = +1 or −1 respectively):

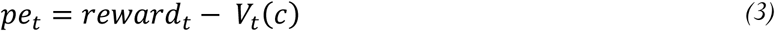

We used two learning rate parameters to model prediction error sensitivity based on the rationale set out previously (Christakou et al., 2013; Bell et al., 2019); the weight of learning from better than expected outcomes (learning rate from positive prediction errors: *η*^*+*^) and the weight of learning from worse than expected outcomes (learning rate from negative prediction errors: *η*^*-*^). The prediction error on each trial was multiplied by either the positive (*η*^*+*^) or negative (*η*^−^) learning rate and used to update the value of the chosen deck:

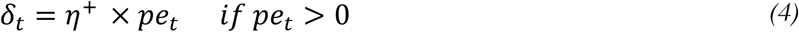

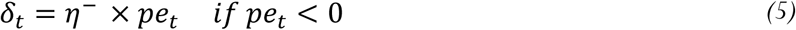

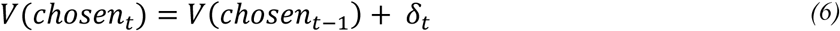

#### Model Fitting

Individuals differ in the degree to which they learn from prediction errors following better-compared to worse-than-expected outcomes (described as learning rate asymmetry), which in turn can affect performance. Generally, it is assumed that the learning rate asymmetry is stable across the learning episode. Reinforcement learning relies on a trade-off between exploration of the available options and exploitation of the optimal choice, which in turn is likely driven by different learning rates. For example, during initial learning, participants must explore the available options to identify the optimal one, therefore they should learn from positive and negative prediction errors equally. However, during periods of stability, and once participants have identified the optimal choice, they must ignore minor losses, placing more weight on positive than negative prediction errors. Indeed, there is evidence that agents able to flexibly alter learning rate asymmetry based on reward history perform better on probabilistic tasks (Cazé and van der Meer, 2013).

To directly test this, we fit the model separately for each task phase (Figure 1B) per participant, estimating parameters that maximised the likelihood of the observed choices given the model (individual maximum likelihood fit; (Daw, 2011)). The calculated deck values from the end of each phase were used as the initial deck values for the following phase, e.g. deck values at the end of the initial learning phase were used as the initial deck values in the first stability phase (Figure 1B). There was no difference in the goodness of fit of the model between task phases when accounting for phase differences in trial number (likelihood repeated measures test: F(3)=0.530, *p=0.664*, partial eta squared= 0.030) (Leong and Niv, 2013; Niv et al., 2015), and therefore parameter estimates from different task phases could be compared. To ensure the model produced consistent, interpretable parameter estimates, *η*^+/-^ was limited to vales between 0 and 1, and *β* and *η*^+/-^ were constrained by gamma- and beta-distributed priors with the following parameters (Christakou et al., 2013):

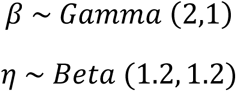

### Functional Magnetic Resonance Imaging

#### Data Acquisition

Data were collected using a Siemens Trio 3T MRI scanner with a 32-channel array head coil at the Centre for Integrative Neuroscience and Neurodynamics (CINN), University of Reading. A multi-band echo planar imaging (EPI) sequence was used to acquire data during the learning task (voxel resolution = 1.8×1.8×1.8 mm; interleaved acquisition of 60 axial slices; no slice gaps; matrix size = 128×128, TE/TR = 40/810 ms; flip angle 31°; multiband factor 6; partial Fourier factor = 1; bandwidth = 1502 Hz/Pixel). This was followed by the acquisition of a high-resolution whole brain T1-weighted structural image using an MPRAGE sequence oriented parallel to the anterior-posterior commissure (voxel resolution = 1×1×1 mm, field of view = 250 mm, 192 sagittal slices, TE/TR = 2.9/2020 ms, flip angle = 9°).

#### Analysis of Functional Data

Analysis was performed using FSL version 5.0.8 (Smith et al., 2004; Jenkinson et al., 2012). First, the brain was extracted from the T1-weighted structural scan using the brain extraction tool (BET) (Smith, 2002). The following pre-statistics processing was used on the functional data: registration to the brain extracted structural scans, followed by linear registration to 1 mm MNI space using FLIRT (Jenkinson and Smith, 2001; Jenkinson et al., 2002); motion correction using MCFLIRT (Jenkinson et al., 2002); brain extraction using BET (Smith, 2002); spatial smoothing (FWHM 3.0 mm); local Gaussian-weighted highpass temporal filtering.

##### Analysis of Cortico- and Thalamo-striatal Interactions

Our main motivation was to dissociate the contribution of lateral OFC and CM-Pf connectivity with the striatum in aspects of reversal learning as described in the introduction. Medial OFC-striatal connectivity was used as an active control, given substantial evidence of medial OFC involvement in reinforcement learning (Morris et al., 2016). Similarly, the MD thalamus also projects to the striatum (Haber and Calzavara, 2009), but (unlike the CM-Pf) it does not project to the striatal cholinergic interneurons (Gonzales and Smith, 2015), which are thought to be recruited by CM-Pf–dorsal-striatal connections during reversal learning (Bradfield et al., 2013; Bell et al., 2018). Therefore, MD-striatal connectivity was also used as an active control. In humans, coarse neuroimaging indiscriminately implicates the thalamus as a whole in behavioural flexibility (Dombrovski et al., 2015; Liu et al., 2015; Shiner et al., 2015). By contrast, our approach here takes into account animal evidence, where MD lesions specifically increase perseveration (Chudasama et al., 2001), whereas CM-Pf lesions increase regressive errors (Bradfield et al., 2013).

For each participant, seed ROI masks were generated for four areas: medial OFC, lateral OFC, MD, and CM-Pf. Separate masks were generated for left and right ROIs, resulting in eight masks in total.

The medial OFC mask consisted of a rectangular box (14 × 22 × 2 mm^3^) centered on the MNI coordinates (x,y,z) ±6, 36, −22. The lateral OFC mask consisted of a rectangular box (6 x 26 x 4 mm^3^) centered on the MNI coordinates ±6, 36, −22 (Morris et al., 2016).

Thalamus masks were generated based on the co-ordinates from Metzger et al. (2010), who used a behavioural task designed to separate thalamic activation in relation to emotional arousal (mediodorsal thalamus) versus attention and expectancy (CM-Pf). Based on this work, we created 6 mm spherical ROIs surrounding the peak voxel in each thalamic region (MNI coordinates (x,y,z): Mediodorsal: ±9.00,-17.00, 14.00; CM-Pf: ±3.78,-16.92, 0.22; Figure 2). The activation time-series were extracted from each ROI and used as parametric modulators in general linear models (GLM) to perform PPI analysis.

**Figure 2.**
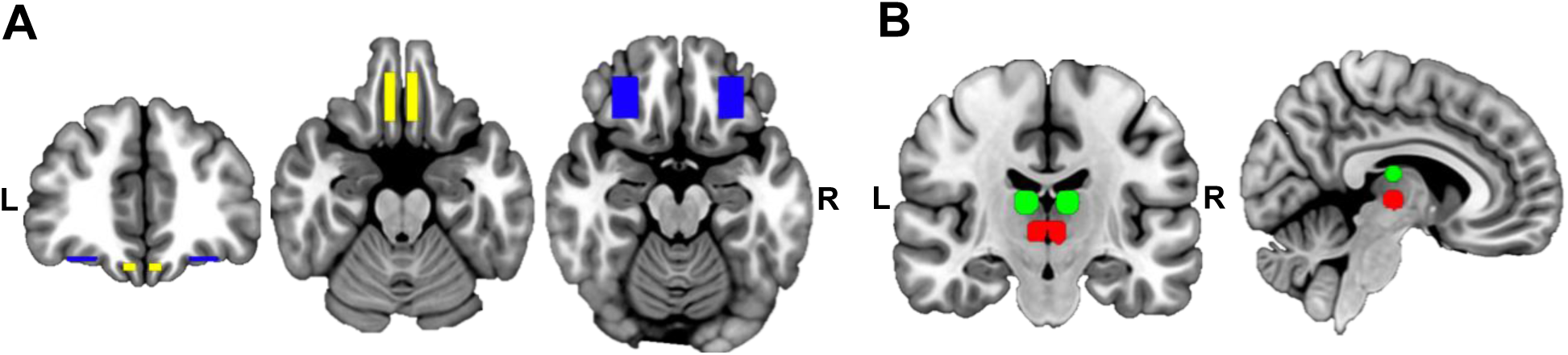
Regions of interest. A. Location of the medial orbitofrontal cortex (mOFC; yellow; MNI coordinates: ±6, 36, −22; number of voxels = 624) and lateral orbitofrontal cortex (lOFC; blue; MNI coordinates: ±28, 36, −18; number of voxels = 616) B. Location of the medio-dorsal (MD; green; MNI coordinates: ±9.00,-17.00, 14.00; number of voxels = 925) and centro-median parafascicular complex (CM-Pf; red; MNI coordinates: ±3.78,-16.92, 0.22; number of voxels = 766) thalamic masks.

**Figure 3.**
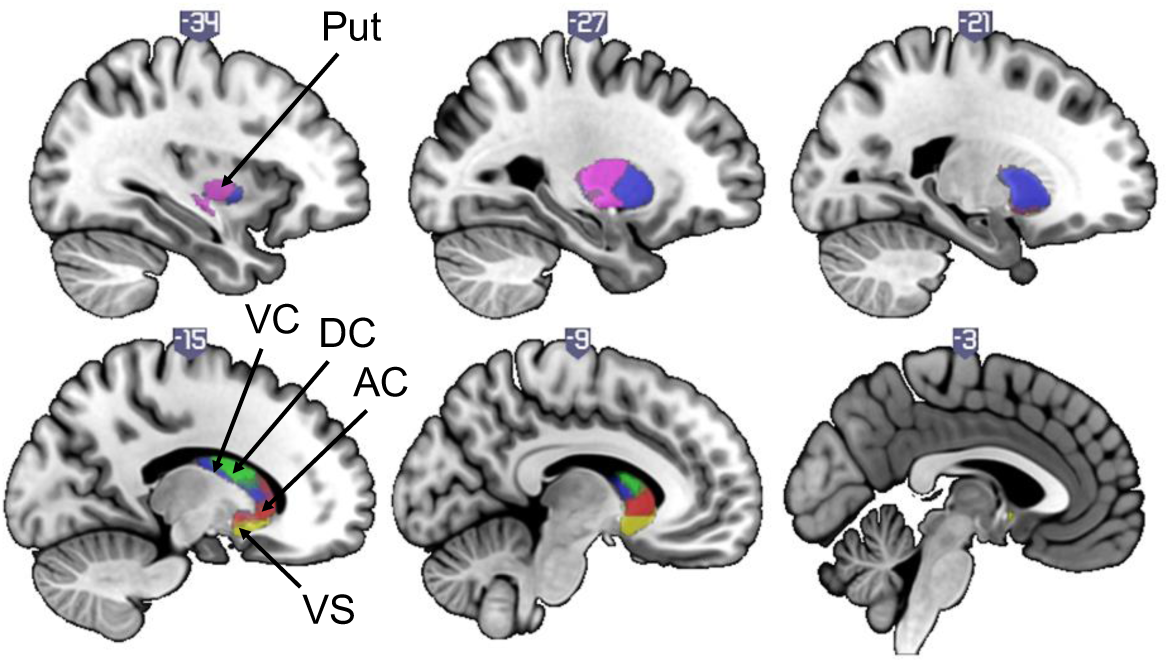
Striatal functional subdivisions. Location and extent of the masks for the dorsal/associative striatum (comprising of anterior caudate (AC; red; MNI coordinates: −12.37, 14.90, −0.27; number of voxels = 2067), dorsal caudate (DC; green; MNI coordinates: −13.51, 6.12, 14.61, number of voxels = 1607), ventral caudate (VC; blue; MNI coordinates: −22.01, 6.66, 2.07; number of voxels = 5208)), putamen (Put; purple; MNI coordinates: −28.90, −8.89, 2.57; number of voxels =2597), and ventral striatum (VS; yellow; MNI coordinates: −11.21, 11.08, −8.45; number of voxels =1219).

A general linear model (GLM) was generated which included four regressors based on the timings for each task epoch (Figure 1A). The regressors were created by convolving a box car function representing the onset and duration of an epoch with an ideal haemodynamic response function. Additionally, the feedback epoch was also modulated by including the prediction error for each trial as generated by the TDRL model as a parametric modulator. Positive and negative prediction errors were included in separate regressors. The ROI timeseries were also included in the GLM, resulting in seven regressors. A separate GLM was created for each area, resulting in eight GLMs in total (left and right lateral OFC, medial OFC, CM-Pf, and MD). The FEAT tool was used to create interaction regressors between the ROI timeseries and the decision making and feedback epochs (Figure 1A; not modulated by prediction error). Higher level analysis was used to generate a group average and contrasts between initial learning and reversal learning. Age was included as a covariate as several brain regions are still maturing in the age range of the sample (Waegeman et al., 2014). Additionally, the number of trials in each phase was included as a covariate to control for different sized data sets, because of individual and task-phase differences in reaching performance criterion.

Striatal target ROI analyses enabled us to focus on the corticostriatal and thalamostriatal connectivity dyads of interest, and were carried out using cluster thresholding (z=2.3, p < 0.05). Striatal target ROIs were defined using five functional subdivisions based on corticostriatal connectivity patterns shown in Choi et al., 2012. The three associative subdivisions in the caudate, extending into the anterior putamen, were summed for ROI analysis of the associative or dorsal striatum. Additionally, a motor ROI was defined in the posterior putamen, and a limbic ROI in the ventral striatum.

### Experimental Design and Statistical Analysis

Statistical analysis was performed using SPSS (IBM Corp. Released 2013. IBM SPSS Statistics for Windows, Version 23.0. Armonk, NY: IBM Corp). Repeated-measures ANOVA tests were used to investigate changes in model parameters across task phases, and to illustrate the PPI analysis results (measures of connectivity were extracted and compared across task phases). When assumptions of sphericity were violated, the Greenhouse-Geisser correction was used. Post hoc tests were conducted using Bonferroni-corrected pairwise comparisons of marginal means. Regression was used to assess interaction effects between brain connectivity, model parameters, and behaviour (mediation analysis, Hayes and Little, 2017). When assumptions of normality were not met (assessed with the Kolmogorov-Smirnov test, Corder & Foreman, 2014) we used bootstrapping (over 1000 samples) to assess the distribution of analysis coefficients (with 95% confidence intervals), and relevant variables were square root-transformed for presentation. Where reporting correlation coefficients is helpful, we used non-parametric analysis (Kendall’s *tau-b*) with bootstrapping.

## RESULTS

### Task Performance

Twenty two (22) participants reached the a priori performance criterion both during initial learning and after the reversal (see Table 1 for trial numbers per task phase).

**Table 1.** Average number of trials per task phase (N=22, SD: standard deviation)

|  | Average Number of Trials | SD |
| --- | --- | --- |
| Initial Learning (to criterion) | 44 | 22 |
| First Stability Phase | 30 | 14 |
| Reversal Learning (to criterion) | 40 | 21 |
| Perseverative errors | 9 | 10 |
| Regressive errors | 6 | 11 |
| Second Stability Phase | 27 | 14 |
| Total | 141 | 58 |

We used a simple reinforcement learning computational model which parameterises aspects of performance that potentially contribute differentially to initial compared to reversal learning, and, further, may have a dissociable effect during performance at criterion. To test this, we compared model parameter estimates across the four task phases (Figure 1B; namely initial learning, performance at criterion after initial learning (first stability period), reversal learning, performance at criterion after reversal learning (second stability period)). The model parameters capture the impact of the subjective value of decisions (value impact parameter, *β*), and the learning rate from positive or negative prediction errors (*η*^*+*^ and *η*^*-*^ respectively).

#### The impact of subjective value increases across the learning episode

Figure 4 (columns) shows that the impact of the subjective value on decisions (*β*) increased over time across the four phases. Specifically, there was a significant effect of task phase on the value impact parameter (β; F(1.8,36.8) = 6.236, p = 0.006, Greenhouse-Geisser corrected, partial eta squared = 0.229). Bonferroni-corrected *post hoc* tests showed that the β value during initial learning was significantly lower than the β values of all other task phases (first stability phase, p = 0.004; reversal learning phase, p = 0.003; second stability phase, p = 0.013; Figure 4).

**Figure 4.**
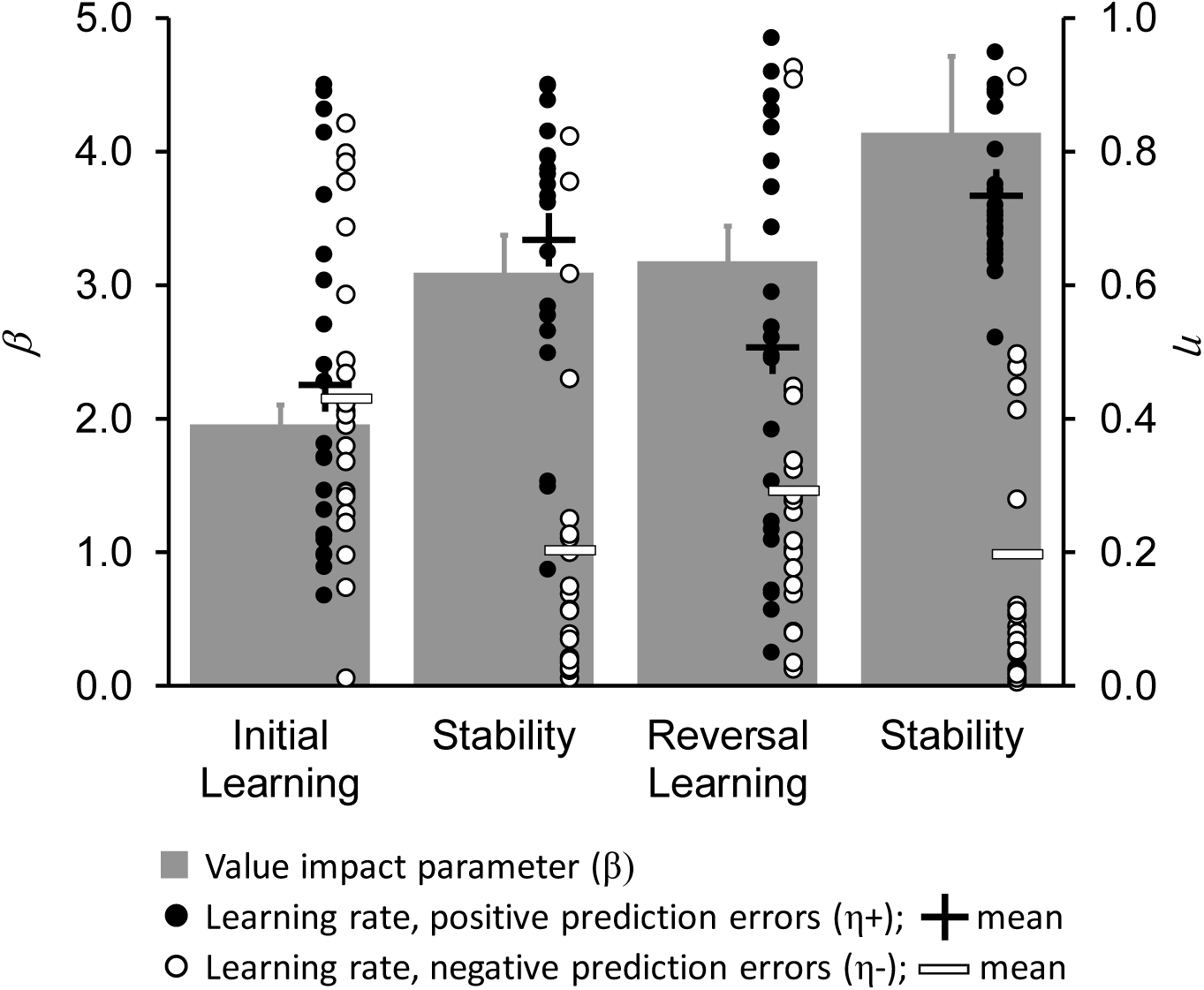
Behavioural model parameter estimates across the task phases. Model parameter estimates changed across the four task phases. The value impact (inverse temperature) parameter (β; columns with error bars denoting the standard error) progressively increased, reaching maximum value during the post-reversal stability period. The relative impact of positive and negative prediction errors can be tracked in the changes in the η+ and η-learning rates (described in the text; η+ black dots with plus signs denoting sample means; η-white dots with minus signs denoting sample means). Learning rates were roughly equal across the sample during initial learning, and changed to favour positive prediction errors during the first stability period. Reversal learning saw both a reduction in η+ and an increase in η-, before recovering after performance criterion was reached once more during the second stability phase.

#### Reversal learning is associated with a return to attending to worse than expected outcomes

Figure 4 (dots) shows that the relative impact of positive and negative prediction errors on choices changed across task phases (described in detail below): after initial learning, participants relied more on positive prediction errors, suppressing the impact of negative prediction errors. But when the contingencies reversed, they reverted to weighing positive and negative prediction errors more equally until they learned to criterion, after which the learning rates diverged again.

Specifically, there was a significant effect of task phase on both learning rates (learning rate from positive prediction errors, η^+^; F(3,63) = 7.021, p < 0.001, partial eta squared = 0.251; learning rate from negative prediction errors, η^-^; F(3,63) = 7.091, p < 0.001, partial eta squared = 0.252). Post hoc tests using the Bonferroni correction revealed η^+^ differed significantly between the initial learning phase and the second stability phase (p = 0.001) and between the reversal learning phase and the second stability phase (p = 0.031). η^+^ did not differ significantly between the other task phases. η^-^ differed significantly between the initial learning phase and the first stability phase (p = 0.003) and between the initial learning phase and the second stability phase (p = 0.004). η^-^ did not differ significantly between the other task phases.

### Analysis of Corticostriatal and Thalamostriatal Interactions

We used seed regions of interest in the medial and lateral OFC, the CM-Pf, and the MD to perform psychophysiological interaction analyses within the three striatal functional subdivisions (dorsal and ventral striatum, and putamen).

This analysis was designed to test the hypothesis that different thalamostriatal dyads would show different connectivity changes across the task phases. Specifically, we tested whether the CM-Pf would show increased connectivity with dorsal associative striatal regions during reversal but not during initial learning, as is the case in the rodent brain (Bradfield et al., 2013). We also asked whether medial and lateral OFC subregions would be associated with initial and reversal learning respectively, as previously suggested.

Finally, we aimed to discriminate the predicted contributions to reversal learning of lateral OFC and CM-Pf connectivity with the striatum, by examining the association of its strength with performance and model parameters.

#### Medial and lateral OFC connectivity with the striatum contribute to initial and reversal learning respectively

PPI analysis revealed a significant correlation between feedback epoch activity in the left medial OFC and activity in the left ventral striatum during the first stability phase (Figure 5A and B), but no other task phase.

**Figure 5.**
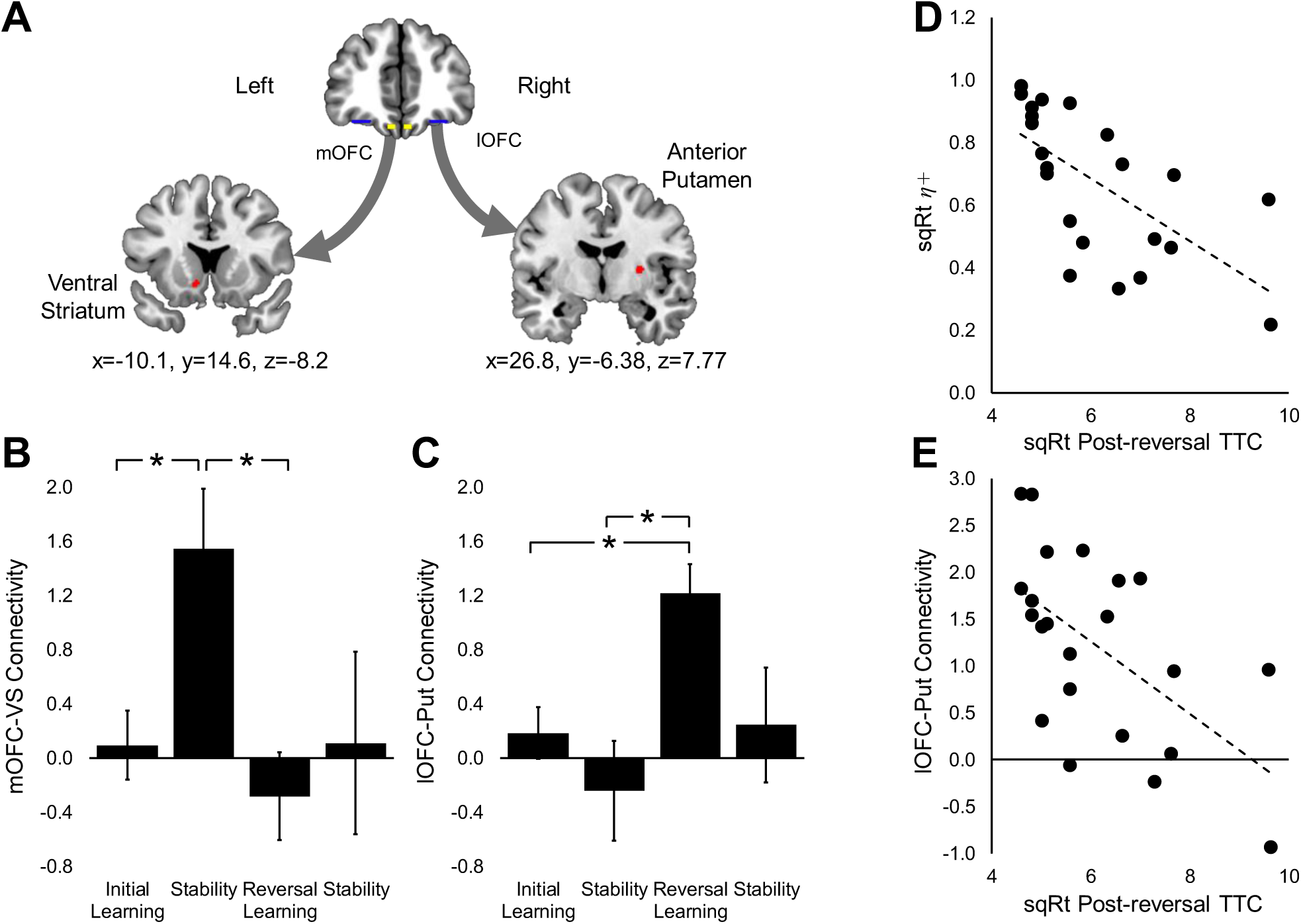
Corticostriatal connectivity and its relationship with performance and model parameters. A. Group average map of the PPI analysis results showing a significant correlation between activation in the left medial OFC and the ventral striatum (SVC p < 0.05; MNI coordinates centre: x = −10.1, y = 14.6, z = −8.2; number of voxels = 58; max Z = 3.45) during the feedback epoch in the first stability phase, and a significant correlation between activation in the right lateral OFC and the anterior putamen (SVC p < 0.01; MNI coordinates centre: x = 26.8, y = −6.38, z = 7.77; number of voxels = 127; max Z = 3.12) during the feedback epoch in the reversal phase. B. Significant effect of task phase on connectivity between the left medial OFC and the left ventral striatum (F(2.14, 45.02) = 3.858, p = 0.026, Greenhouse-Geisser corrected, partial eta squared = 0.155). Connectivity was significantly higher during the first stability phase compared to all other task phases. C. Significant effect of task phase on connectivity between the right lateral OFC and the right anterior putamen (F(2.26, 47.55) = 3.891, p = 0.023, Greenhouse-Geisser corrected, partial eta squared = 0.156). Connectivity was significantly higher during the reversal phase compared to all other task phases. D and E. Bivariate correlations between post-reversal trials to criterion and η+ (panel D; Kendall’s tau-b = −0.592, p < 0.001), and between lateral OFC-putamen connectivity and trials to criterion in the reversal learning phase (panel E; Kendall’s tau-b = −0.415, p = 0.008). mOFC: medial orbitofrontal cortex; lOFC: lateral orbitofrontal cortex; VS: ventral striatum; Put: putamen; error bars represent the standard error; *p < 0.05.

To illustrate this effect, we extracted the average estimate of connectivity in the significant cluster for each task phase: a repeated-measures ANOVA showed a significant main effect of task phase on connectivity (F(2.14, 45.02) = 3.858, p = 0.026, Greenhouse-Geisser corrected, partial eta squared = 0.155). Medial OFC-ventral striatal connectivity was higher during the first stability phase compared to other task phases (Figure 5B): initial learning: mean difference = 1.452, p=0.037; reversal learning: mean difference = 1.829, p = 0.016; second stability period: mean difference = 1.437, p = 0.408.

We also observed a significant correlation between feedback epoch activity in the right lateral OFC and activity in the right anterior putamen during the reversal phase (Figure 5A and C). There were no significant correlations between lateral OFC and striatal activity during initial learning, nor during the stability phases. Reflecting the neuroimaging analysis, repeated-measures ANOVA showed a significant main effect of task phase on connectivity (F(2.26, 47.55) = 3.891, p = 0.023, Greenhouse-Geisser corrected, partial eta squared = 0.156). Lateral OFC-dorsal striatal connectivity was higher during the reversal learning phase compared to other task phases (Figure 5C): initial learning: mean difference = 1.035, p=0.002; first stability phase: mean difference = 1.461, p = 0.010; second stability phase: mean difference = 0.976, p = 0.399.

#### CM-Pf connectivity with the dorsal striatum increases specifically during reversal learning

During reversal learning, PPI analysis revealed a significant correlation between feedback epoch activity in the left CM-Pf and activity in the left dorsal striatum (Figure 6A and C) (corresponding to the ventral caudate (VC) subregion as defined in Choi et al. 2012, see Figure 3). There were no significant correlations between CM-Pf and dorsal striatum activity during initial learning, nor during the stability phases. This connectivity finding was specific to the CM-Pf, with no significant correlations between activation in the MD and the dorsal striatum.

**Figure 6.**
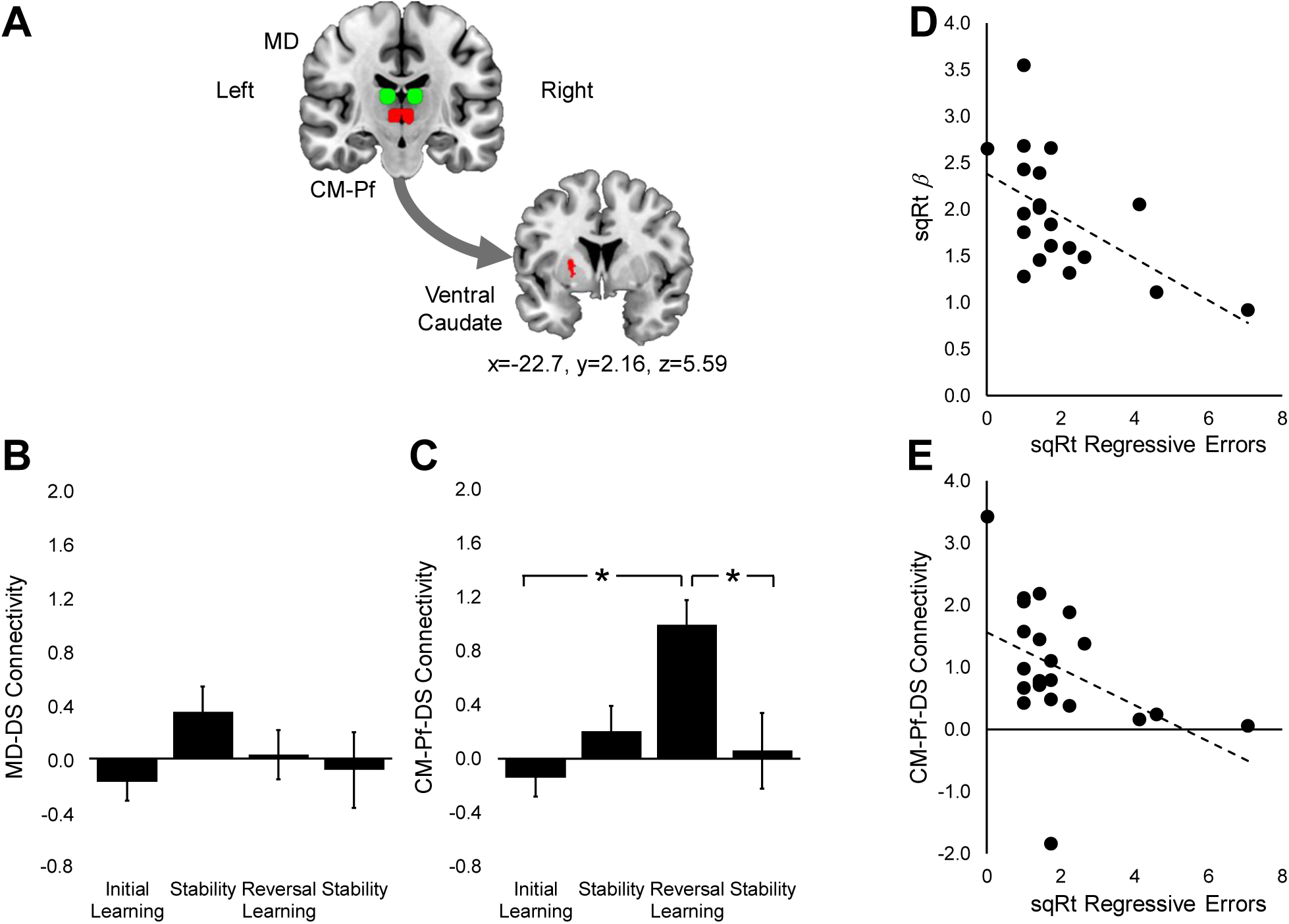
Thalamostriatal connectivity and its relationship with performance and model parameters. A. Group average map of the PPI analysis results showing a significant correlation learning between activation in the left dorsal striatum (specifically ventral caudate) and the left CM-Pf (SVC p < 0.05; MNI coordinates centre: x = −22.7, y = 2.16, z = 5.59; number of voxels = 151; max Z = 3.01) during the feedback epoch in the reversal phase. B. There was no significant effect of task phase on connectivity between the left MD and the dorsal striatum (F(3,63) = 1.145, p = 0.338, partial eta squared = 0.052). C. Significant effect of task phase on connectivity between the left CM-Pf and the left dorsal striatum (F(3,63) = 5.510, p = 0.002, partial eta squared = 0.208). Connectivity was significantly higher during the reversal learning phase compared to all other task phases. D and E. Bivariate correlations between regressive errors and CM-Pf – dorsal striatum connectivity during reversal learning (panel D; Kendall’s tau-b = −0.415, p = 0.010), and between regressive errors and the value impact parameter (β) during the post-reversal stability period (panel E; Kendall’s tau-b=0.333, p=0.030). CM-Pf: centro-median parafascicular thalamic complex; DS: dorsal striatum; MD: mediodorsal thalamus; error bars represent the standard error; *p < 0.05.

To illustrate the effect, we extracted the average strength of CM-Pf–dorsal-striatal connectivity for each task phase: a repeated-measures ANOVA showed a significant main effect of task phase on connectivity (F(3,63) = 5.510, p = 0.002, partial eta squared = 0.208). Connectivity was higher during reversal learning compared to the other task phases (Figure 6C): initial learning phase: mean difference = 1.135, p = 0.015; first stability phase: mean difference = 0.788, p=0.091; final stability phase: mean difference = 0.934, p=0.033.

By contrast, reflecting the lack of effect in the ROI PPI analysis, there was no significant effect of task phase on connectivity between the left MD and dorsal striatum (F(3,63) = 1.145, p = 0.338, partial eta squared = 0.052; Figure 6B).

### Distinct Neurocognitive Mechanisms of Reversal Learning

#### lOFC-striatal connectivity speeds up reversal by increasing the learning rate from positive prediction errors

Figure 5E and Table 2 show that increased lateral OFC-striatal connectivity during reversal learning was associated with reduced trials to reach criterion.

**Table 2.**
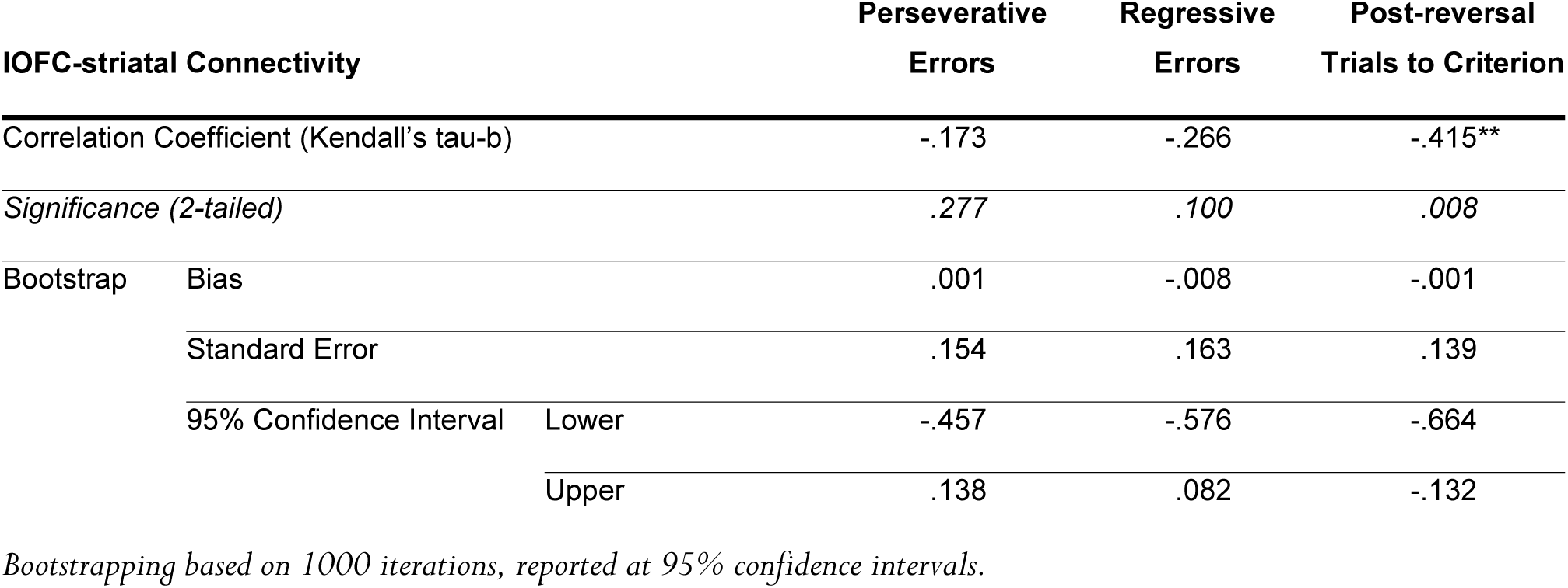
Strength of corticostriatal connectivity is associated with speed of reversal

The only model parameter that also correlated with speed of learning (trials to criterion) in either initial or reversal learning was η+ during reversal learning, showing a strong negative correlation with trials to criterion (Kendall’s tau-b =-0.592, p<0.001; 1000 iteration bootstrap: bias = 0.001, 95% CI: −0.373 to −0.772; shown in Figure 5D). Figure 4 shows that η-increased slightly across the sample during the reversal phase, as participants started to encounter unanticipated losses. By contrast, although the mean of η+ decreased, its variance increased. Given this profile, the correlation with trials to criterion may simply describe the fact that participants who learn the reversal faster return to encountering positive outcomes more quickly. At the same time, however, it is possible that η+ remains high in participants who have a better task representation (e.g. understand that there are better and worse options available at any given time), which would in turn accelerate learning.

To explore the mechanistic relationship of these variables, we tested for a mediating influence of η+ on the relationship between lateral OFC-striatal connectivity and speed of reversal, using bootstrapping (Hayes and Little, 2017; Hayes and Rockwood, 2017). The conditional model is described in Table 3, including the underlying correlation assumptions. This analysis suggests that if the lateral OFC-striatal connectivity observed during reversal leads to faster reversl learning, it may do so through a mechanism that helps maintain a high η+, despite the occurrence of unexpected negative outcomes during this phase.

**Table 3.**
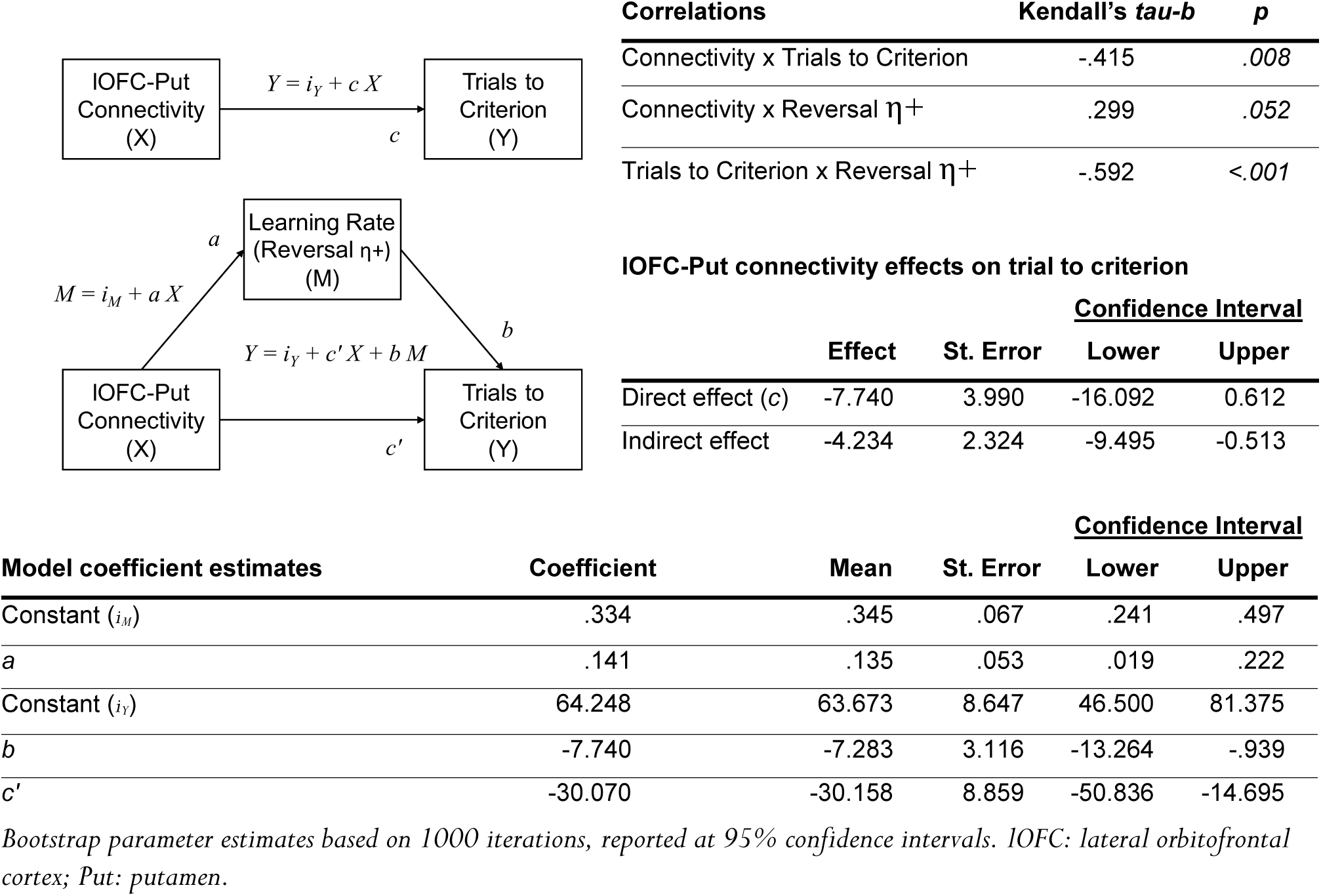
The learning rate from positive prediction error (η+) mediates the relationship between lateral OFC-striatal connectivity and speed of reversal

| Model coefficient estimates | Coefficient | Mean | St. Error | Confidence Interval |  |
| --- | --- | --- | --- | --- | --- |
|  |  |  |  | Lower | Upper |
| Constant ($i_M$) | .334 | .345 | .067 | .241 | .497 |
| *a* | .141 | .135 | .053 | .019 | .222 |
| Constant ($i_Y$) | 64.248 | 63.673 | 8.647 | 46.500 | 81.375 |
| *b* | -7.740 | -7.283 | 3.116 | -13.264 | -.939 |
| *c'* | -30.070 | -30.158 | 8.859 | -50.836 | -14.695 |

Bootstrap parameter estimates based on 1000 iterations, reported at 95% confidence intervals. IOFC: lateral orbitofrontal cortex; Put: putamen.

#### CM-Pf-striatal connectivity prevents regressive errors by promoting new learning within an established task representation

There is evidence in the animal literature that disruption of CM-Pf–dorsal striatal connections specifically increases regressive (as opposed to perseverative) errors after reversal (Bradfield et al., 2013; Bradfield and Balleine, 2017). In line with this evidence, increased CM-Pf-dorsal-striatal connectivity during reversal was associated specifically with reduced regressive errors (Figure 6E and Table 4) (but not with perseverative errors), even though it had no effect on the overall speed of criterion learning (Table 4).

**Table 4.**
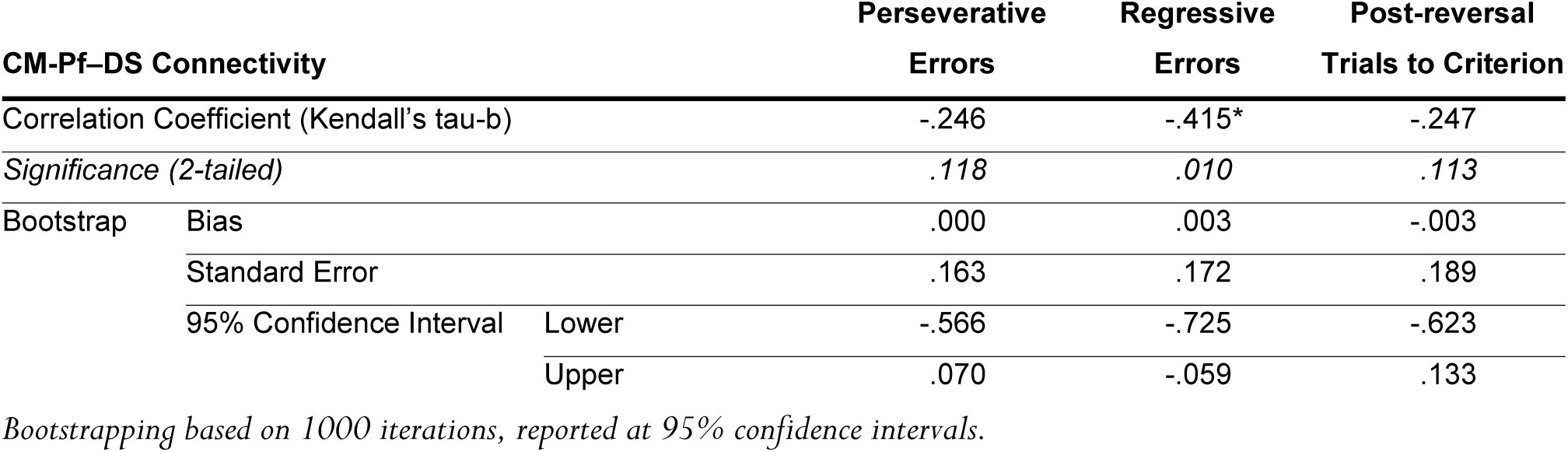
Strength of thalamostriatal connectivity is associated with regressive errors

The only model parameter to also show a negative correlation with regressive errors was the value impact parameter β of the post-reversal stability period (Kendall’s tau-b =-0.443, p=0.006; 1000 iteration bootstrap: bias = −0.009, 95% CI: −0.751 to −0.107; shown in Figure 6D).

Similar to the OFC analysis in the previous section, in order to explore the mechanistic relationship between these variables we performed a mediation analysis with bootstrapping (Hayes and Little, 2017; Hayes and Rockwood, 2017), to describe the indirect effect of CM-Pf dorsal striatal connectivity on regressive errors via the mediating effect of post-reversal β. The conditional model is described in Table 5, including the underlying correlation assumptions. The analysis suggests that the effect of CM-Pf-dorsal-striatal connectivity in reducing regressive errors is mediated (completely, in this sample) by a mechanism that promotes new contingency learning within a more stable longer-term task representation, and is read-out in the increased impact of value history on post-reversal decisions.

**Table 5.**
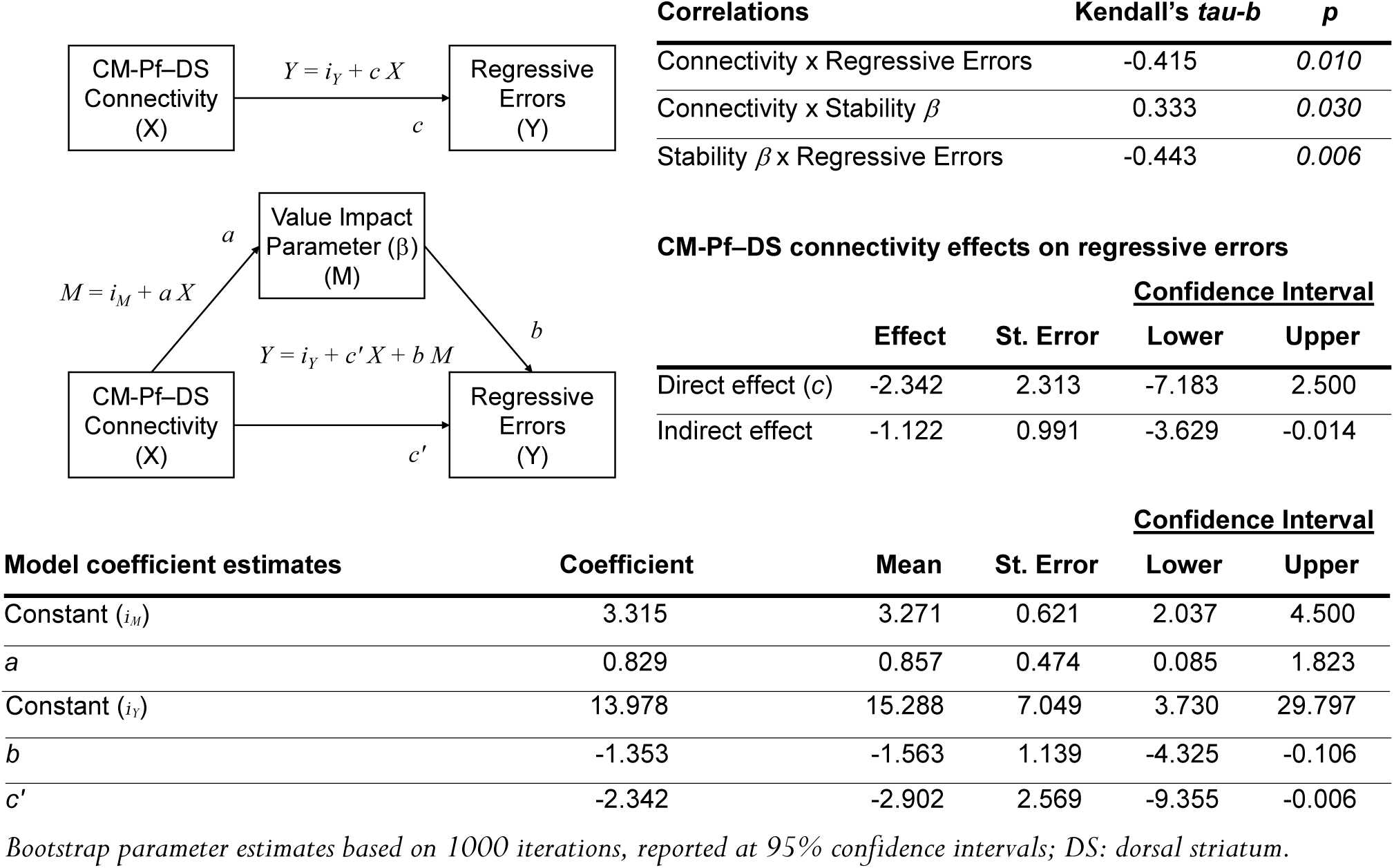
Inverse temperature (β) mediates the negative correlation between CM-Pf–dorsal striatal connectivity and regressive errors

## DISCUSSION

We investigated the role of corticostriatal and thalamostriatal connectivity in human reversal learning by combining functional connectivity analysis of fMRI data and computational modelling. We demonstrate, for the first time in humans, evidence for the role of connectivity between CM-Pf and dorsal striatum in reversal learning. In line with animal experiments, the strength of this connectivity was associated with reduced regressive errors, a key component of effective probabilistic reversal. In addition, in line with both the human and animal literature, we show specific roles for medial and lateral OFC connectivity with the striatum, during initial and reversal learning respectively.

Specifically, functional connectivity between the CM-Pf and the dorsal striatum was observed only during reversal learning, in line with evidence from the animal literature. In rodents, disrupting connectivity between the CM-Pf and the dorsomedial (associative) striatum results in impaired reversal learning, whilst initial learning remains intact. The impact of such a manipulation on reversal learning is specific: animals are able to identify the reversal, but display interference from the initial learning when learning and expressing the new behaviour. This effect is thought to be driven by disruption of input specifically to the cholinergic system in the striatum (Brown et al., 2010; Bradfield et al., 2013).

The connectivity effect described above was specific to the CM-Pf. There was no significant correlation between activity in the mediodorsal thalamus and the dorsal striatum during reversal, and no significant change in connectivity between these regions across any task phases, which serves as an important functional control: the mediodorsal thalamus also projects to the dorsal striatum (Haber and Calzavara, 2009), but it does not project to the striatal cholinergic system (Gonzales and Smith, 2015), which is thought to interface the CM-Pf influence into corticostriatal function (Smith et al., 2014). Indeed, in a study in humans with the same task presented here, we recently showed evidence of cholinergic recruitment in the same dorsal striatal region during reversal learning using functional magnetic resonance spectroscopy (fMRS) (Bell et al., 2018). Together, these observations are in line with the notion that reversal-specific changes in CM-Pf-dorsal striatal connectivity relate to changes in recruitment of the striatal cholinergic system.

Connectivity between the lateral OFC and the anterior putamen was also increased during the reversal learning period, in line with previous evidence. For example, inactivation of the lateral OFC reduces win-stay/lose-shift behaviour in rats only after incorrect choices, and is therefore thought to adjust response selection following violation of reward expectancies (Dalton et al., 2016).

Functional connectivity between the medial OFC and the ventral striatum was significant after initial learning, once performance had reached criterion, and before the reversal was implemented. The medial OFC is required for computing value representations, and inactivation of the medial OFC has been shown to impact both initial and reversal learning, possibly due to a reduction in the ability to incorporate positive or negative feedback to guide action selection (Dalton et al., 2016). Medial OFC activation in humans reportedly discriminates stimuli to which responses need to be inhibited following contingency reversal, with response strength predicting behavioural learning strength (Zhang et al., 2015). Further, Morris et al. (2016) showed that a measure of model-based behaviour (or goal-directedness) positively correlated with connectivity between the medial OFC and ventral striatum. In the task reported here, during the first stability period (i.e. after reaching criterion), participants should have formed a reliable model of the task and successful participants use this model to guide their actions, ignoring the occasional negative feedback, as indeed indicated by a decrease in the learning rate for negative prediction errors, η-.

We observed changes in learning rate asymmetry (the relative impact of learning rates from positive and negative prediction errors) throughout the task. It has been shown previously that agents able to flexibly alter learning rate asymmetry based on reward history perform better on a probabilistic task (Cazé and van der Meer, 2013). However, the contribution of learning rate asymmetry to different stages of learning during probabilistic reversal remains unclear (Krugel et al., 2009; Javadi et al., 2014). Our simple reinforcement learning model enabled us to track changes in the relative influence of positive and negative prediction errors at different task phases. During initial learning, participants place equal weights on positive and negative prediction errors to identify which decks provide overall wins and losses. By the first stability period, after criterion has been reached, participants have identified the optimal deck and are able to ignore any probabilistic losses. Consequently, we observed increased learning rate from positive prediction errors and decreased learning rate from negative prediction errors. During the reversal learning phase, participants start receiving more negative feedback. If they are to identify that this is no longer experienced as part of the learned probabilistic structure, but rather that the contingencies have changed, participants must increase the weight of learning from negative prediction errors, as was observed. This way, attending to worse than expected outcomes provides the opportunity to adaptively dismantle confidence in the previously learned response, making the change in the relative learning rates an important marker of reversal learning efficiency. During the last stability period, participants will have again identified the optimal deck and may once again ignore any probabilistic losses, recovering the divergence in learning rates. Therefore, by continuously updating the learning rates based on feedback, participants are able to adapt to alterations in task structure. This is an important component of reversal learning, and cognitive flexibility more generally, and provides an insight into the more nuanced skills required for the ability to flexibly alter behaviour. Using this framework, we expand the evidence for OFC involvement in task set representation to OFC-striatal connectivity: η+ correlated with lOFC-striatal connectivity, and further mediated its relationship with the speed of reversal (indexed by the number of trials to reach post-reversal criterion).

Changes in the relative learning rates from positive and negative prediction errors were accompanied by increases in the impact of subjective value on choice behaviour, an effect that is uncontroversial in simple probabilistic learning tasks (Krugel et al., 2009). But here we also show that the value impact parameter β continues to increase after the reversal is overcome, suggesting a cumulative learning effect which we speculate may be related to task-structure learning rather than stimulus-outcome learning (e.g. that identifying the highest yielding option reduces all the uncertainty in the task).

Finally, we show evidence that the impact of subjective value post-reversal may mediate the relationship between the strength of the CM-Pf-dorsal striatal connectivity and successful adaptation in the task. Importantly, stronger connectivity was associated specifically with fewer regressive errors, while there was no association between connectivity and perseveration. This is in line with evidence from the animal literature, where disrupting CM-Pf-dorsal striatal connectivity results in an increase in the number of regressive errors, whilst there is no effect on perseveration (Bradfield et al., 2013). This is thought to represent interference between new and existing learning, resulting from an inefficient partition of the conflicting contingencies into separate internal states (Bradfield and Balleine, 2017). It has been suggested that the initial and reversed contingencies are encoded in separate pools of neurons within the dorsal striatum. CM-Pf controlled cholinergic modulation may be used to select the appropriate pool of neurons for encoding and selecting an action based on the internal state (Stalnaker et al., 2016; Bradfield and Balleine, 2017). This may be achieved by decoupling CIN firing from dopaminergic control to prevent extinction learning, while reseting corticostriatal dynamics to enable new associations to drive behaviour (Ashby and Crossley, 2011).

In summary, we provide evidence for the involvement of CM-Pf–dorsal striatal connectivity in reversal learning, showing its strength to be associated with the efficiency of behavioural adaptation. We also demonstrate the involvement of medial and lateral OFC-striatal connectivity to initial learning and reversal. Finally, applying our reinforcement learning model, we discriminate the contributions of corticostriatal and thalamostriatal systems in reversal learning. The study helps to bridge the gap between animal studies of this system and human studies of reversal learning and cognitive flexibility more generally, and highlights the contribution of thalamostriatal connectivity to these critical competences.

## Acknowledgments

This study was supported by a Human Frontier Science Program (HFSP) grant (RGP0048/2012), and an Engineering and Physical Sciences Research Council (EPSRC) doctoral training grant (EP/L505043/1). The funders had no involvement in study design, in the collection, analysis, and interpretation of data, in writing the report, or in the decision to submit the paper for publication.

